# A Systems Biology Approach to Pathogenesis of Gastric Cancer: Gene Network Modeling and Pathway Analysis

**DOI:** 10.1101/2022.09.12.507635

**Authors:** Negar Mottaghi-Dastjerdi, Abozar Ghorbani, Hamed Montazeri

**Affiliations:** Department of Pharmacognosy and Pharmaceutical Biotechnology, School of Pharmacy, Iran University of Medical Sciences, Tehran, Iran; Nuclear Agriculture Research School, Nuclear Science and Technology Research Institute (NSTRI), Karaj, Iran

**Author notes:** **Corresponding author:** Dr. Negar Mottaghi-Dastjerdi, Department of Pharmacognosy and Pharmaceutical Biotechnology, School of Pharmacy, Iran University of Medical Sciences, Tehran, Iran. **Co-corresponding author:** Dr. Abozar Ghorbani, Nuclear Agriculture Research School, Nuclear Science and Technology Research Institute (NSTRI), Karaj, Iran.

**Keywords:** Biological networks, Differentially expressed genes, Gastric cancer, Gene ontology, Kyoto Encyclopedia of Genes and Genomes, Promoter analysis, Protein-protein interactions

## Abstract

**Introduction:** Gastric cancer (GC) ranks among the most common malignancies worldwide. In our previous study, we found overexpressed genes in GC clinical samples. The goal of the current study was to find critical genes and key pathways involved in the pathogenesis of GC.

**Methods:** Gene interactions were analyzed using STRING, and Cytoscape was used to visualize the molecular interaction network. CytoHubba was used for drawing the PPI network and identifying hub proteins. The Kyoto Encyclopedia of Genes and Genomes (KEGG) and Gene Ontology (GO) at STRING were used for the enrichment analysis of the hub genes. Cluster analysis of the network was done using CytoCluster. MEME Suite was used for promoter analysis of the hub genes using Tomtom and GoMo tools.

**Results and Discussion:** Our results showed that the most affected processes in GC are the metabolic processes. The OXPHOS pathway was also considerably enriched in our analyses. These results showed the significant role of mitochondria in GC pathogenesis. Although many investigations have focused on the mitochondrial role in the pathogenesis of various cancers, the characteristics of respiratory and metabolic changes in GC have not been fully elucidated. Our results also showed that most of the affected pathways in GC were the pathways also involved in neurodegenerative diseases. Also, promoter analysis showed that negative regulation of signal transduction might play an important role in GC pathogenesis. The results of this study might open up new insights into GC pathogenesis. The identified genes might be novel diagnostic or prognostic biomarkers or potential therapeutic targets for GC. Nonetheless, these results were obtained by bioinformatics analysis and require further clinical validation.

## Introduction

Gastric cancer (GC) is the fifth most common cancer and the fourth cause of cancer-related death worldwide, with more than 1 million new cases in 2020 and an estimated 769,000 deaths, equal to one out of each 13 deaths worldwide (1). Although the association between different risk factors and the development of GC has been investigated in different studies (2), the specific molecular network mechanisms have not been fully introduced. Therefore, elucidating the molecular mechanisms involved in GC pathogenesis may help to find targets for early detection and classification and prolong patient survival.

Carcinogenesis is a complicated process involving several genetic and epigenetic alterations (3). Adenocarcinoma is the most common type of GC, which has two subtypes, the diffuse type, and the intestinal type, with different molecular characteristics (4). There are many investigations on the epigenetic and genetic alterations in different types of GC, involving alterations in cell cycle regulators, tumor suppressor genes, DNA repair genes, and oncogenes (5). The exploration of the genes which are dysregulated in different pathways may be beneficial in elucidating the molecular pathophysiological mechanisms underlying carcinogenesis and, therefore, may help to develop new treatment strategies. Recently, gene and network analysis using high-throughput methods have been used as hopeful tools with several clinical applications, including cancer detection and classification, as well as patient response to the treatment and the prognosis of the disease (6, 7). Using systems biology and systems pharmacology approaches can provide a better understanding of mechanisms and detect gene signatures for precision and personalized medicine (8). These types of systems-level insights then can be used to design more accurate in silico models of biological circuits leading to cells and tumor responses (9, 10).

There are many investigations on alterations of the genes in GC. However, they are not still enough to provide an elucidated picture of the molecular pathogenesis of GC. Therefore, along with these studies, and to provide more clues into the mysterious molecular pathogenesis of GC, we previously studied the gene expression profile of GC using the SSH method. We introduced the overexpressed genes in GC (11-13). The present study uses bioinformatics analysis to investigate the genes involved in the pathogenesis of GC introduced in our previous studies and investigates the important GO terms, KEGG pathways, and protein-protein interaction PPI networks, with a specific focus on potential gene hubs which have possible roles in the gastric carcinogenesis.

## Methods

### Selection of the genes involved in gastric carcinogenesis

In our previous studies (11-13), representative overexpressed genes in GC were identified by SSH. In the current study, the genes introduced in these three studies as the overexpressed genes identified in GC were selected for network analysis (**Table 1)**.

**Table 1.** List of the genes used in this study as the overexpressed genes identified in GC by the SSH method (11-13).

| # | Gene name | Gene ID | Cytogenetic location | Description |
| --- | --- | --- | --- | --- |
| 1 | RPL18A | 6142 | 19p13 | 60S ribosomal protein L18a |
| 2 | TSPAN8 | 7103 | 12q14.1–q21.1 | Tetraspanin 8 |
| 3 | COII | 4513 | MT | Cytochrome c oxidase subunit 2 (COII) |
| 4 | ND4 | 4538 | MT | NADH dehydrogenase, subunit 4 |
| 5 | ATP6 | 4508 | MT | Mitochondrially encoded ATP synthase 6 |
| 6 | EIF4A1 | 1973 | 17p13 | Eukaryotic translation initiation factor 4A1 |
| 7 | RNASEH2B | 79621 | 13q14.3 | Ribonuclease H2, subunit B; AicardiGoutieres syndrome 2 protein; RNase H2 subunit B |
| 8 | SEC13 | 6396 | 3p25-p24 | Protein SEC13 homologue |
| 9 | VAT-1 | 10493 | 17q21.31 | Vesicle amine transport protein-1 |
| 10 | *MT-RNR2L1 (HN1)* | 100462977 | 17p11.2 | MT-RNR2 like 1 (pseudogene) |
| 11 | *MT-RNR2L3 (HN3)* | 100462983 | 20q13.31 | MT-RNR2 like 3 (pseudogene) |
| 12 | *MT-RNR2L6 (HN6)* | 100463482 | 7q34 | MT-RNR2 like 6 (pseudogene) |
| 13 | *MT-RNR2L10 (HN10)* | 100463488 | Xp11.21 | MT-RNR2 like 10 (pseudogene) |

### Reconstruction of genes and PPI networks and the hub analysis

STRING is a database of predicted and known interactions of proteins. To evaluate the interactions among the genes listed in Table 1, these genes were analyzed in the web-based application of STRING ver.10 (http://string-db.org) (14) with a minimum required interaction score of 0.15 (low confidence), and the PPI list was prepared. Subsequently, the PPI list was imported into Cytoscape (version 3.9.1), an open-source platform for visualization of molecular interaction networks and data integration. CytoHubba software (Version 0.1) was used to draw the PPI network and identify hub proteins between all nodes (15). Four computational algorithms of Cyto-Hubba, including MCC, Degree, MNCC, and MNC, were used for detecting and ranking the hub genes, and four proteins for each algorithm were selected as hub genes. Subsequently, the hub genes and their interactions were shown in a subnetwork.

### Gene Ontology and pathway enrichment analysis of the hub genes

The enrichment analysis of the hub genes was analyzed based on the Kyoto Encyclopedia of Genes and Genomes (KEGG) and Gene Ontology (GO), including molecular function (MF), cellular components (CC), and biological process (BP) at the web-based application of STRING (14).

### Cluster analysis of the network

CytoCluster (Version 2.1.0) was used for the clustering of network nodes. The IPCA (Identifying Protein Complex Algorithm) algorithm was used for cluster analysis of the subnetwork. This algorithm is density-based, which identifies dense subgraphs in protein interaction networks. IPCA determines the weight of an edge by calculating the common neighbors of its connected nodes and calculates each node’s weight through a sum up of the weights of its incident edges. (16). The threshold was set to 10, and the genes of each cluster were further analyzed in STRING ver.10 (14) to find the KEGG pathways in which these genes are involved.

### Promoter motif analysis of hub genes

The 1 kbp upstream flanking regions (UFRs) of hub genes were extracted from Ensembl BioMart web services (https://asia.ensembl.org/info/data/biomart/index.html). Conserved motifs on the sequences were identified using MEME Suite (version 5.4.1) (meme.nbcr.net/meme/intro.html) (17) with its default parameters except for the threshold P- and E-values of <0.01 and <0.0001, respectively. Tomtom (version 5.4.1) tool (http://meme-suite.org/tools/tomtom) (18) was used to remove redundant motifs and identify known CRE based on the motif database of Human (homo sapiens) DNA (HOCOMOCO Human (v11 Full) with the threshold P- and E-values of <0.01 and <0.0001, respectively. GoMo tool (http://meme-suite.org/tools/gomo) was also used to find potential roles for motifs (19).

## Results and discussion

### Reconstruction of genes and PPI networks and the hub analysis

In the present study, bioinformatic analysis of DEPs was performed in GC based on the identified genes from our previous studies (11-13). The reconstructed gene and PPI networks are shown in Error! Reference source not found.. The hub analysis resulted in the identification of 11 genes with the most interactions (**Table 2)**. The subnetwork prepared by these identified hub genes is illustrated in Error! Reference source not found.. As shown in **Table 2**, four genes encode different subunits of the ATP synthase, including ATP5A1, ATP5B, ATP5D, and MT-ATP8. “F1F0 ATP synthase”, also known as “Complex V” or “mitochondrial membrane ATP synthase”, recruits a proton gradient across the inner membrane, which is generated by electron transport complexes of the respiratory chain to produce ATP from ADP. (20). Liu *et al*. reported ATP5A1 as one of the up-regulated genes in GC (21). ATP5B is one of the most important subunits of ATP synthase and increases cellular ATP levels. Wang *et al*. found that high ATP5B expression in tumor tissues of GC is positively correlated with age, tumor size, the TNM stage, lymph node metastasis, and patients’ poor prognosis. They reported that ATP5B overexpression in GC cells caused ATP-promoting migration, invasion, and proliferation. In their study, an increase in MMP2 expression results in the phosphorylation of FAK, and phosphorylated AKT was observed by ATP5B overexpression in GC cells. Also, an increased level of extracellular ATP happens after ATP5B overexpression through the intracellular ATP secretion, and the FAK/AKT/MMP2 pathway is activated. Activation of the ATP5B-induced downstream pathway is activated through the P2X7 receptor. Inhibition of P2X7, FAK, AKT, and MMP2 results in the suppression of proliferation, migration, and invasion of GC cells. In conclusion, studies by Wang *et al*. showed that ATP5B involves in GC tumor progression through FAK/AKT/MMP2 pathway (22). Therefore, ATP5B may serve as a poor prognosis biomarker and a beneficial therapeutic target for GC. MT-ATP8 is a mitochondrial gene encoding membrane subunit 8 of ATP synthase. Several genetic syndromes that contributed to mitochondrial dysfunction depicted by a reduced OXPHOS ability have been explained (23). MT-ATP8 was reported to have mutations in GC (24). ATP5D encodes the subunit delta of the catalytic core of ATP synthase, F1. Wei *et al*. found that overexpression of PSMB10, VPS13D, NDUFS8, ATP5D, POLR2E, and HADH were correlated with adverse overall survival in acute myeloid leukemia (AML) (25).

**Table 2.** List of hub genes identified by using CytoHubba.

| # | Node | Gene ID | Ensembl | Gene Description | Rank | Ranking Method | Other names | Location |
| --- | --- | --- | --- | --- | --- | --- | --- | --- |
| 1 | ATP5A1 | 498 | ENSG00000152234 | ATP synthase F1 subunit alpha | 1 | MCC, MNC, Degree | OMR; ORM; ATPM; MOM2; ATP5A; hATP1; ATP5F1A; MC5DN4; ATP5AL2; COXPD22; HEL-S-123m | 18q21.1 |
| 2 | ATP5B | 506 | ENSG00000110955 | ATP synthase F1 subunit beta | 2 | MCC, MNC, Degree | ATP5F1B, ATPMB; ATPSB; HEL-S-271 | 12q13.3 |
| 3 | NDUFS3 | 4722 | ENSG00000213619 | NADH: ubiquinone oxidoreductase core subunit S3 | 3 | MCC | CI-30; MC1DN8 | 11p11.2 |
| 4 | COX6C | 1345 | ENSG00000164919 | Cytochrome c oxidase subunit 6C | 4 | MCC | - | 8q22.2 |
| 5 | MT-ND6 | 4541 | ENSG00000198695 | mitochondrially encoded NADH dehydrogenase 6 | 1 | DMNC | MTND6; ND6 | MT (non-nuclear) |
| 6 | MT-ATP8 | 4509 | ENSG00000228253 | mitochondrially encoded ATP synthase 8 | 2 | DMNC | ATPase8; MTATP8; ATP8 | MT (non-nuclear) |
| 7 | MT-ND4 | 4538 | ENSG00000198886 | mitochondrially encoded NADH dehydrogenase 4 | 3 | DMNC | MTND4; ND4 | MT (non-nuclear) |
| 8 | COX7A2 | 1347 | ENSG00000112695 | cytochrome c oxidase subunit 7A2 | 4 | DMNC | VIIAL; COX7AL; COX7AL1; COXVIIAL; COXVIIa-L | 6q14.1 |
| 9 | RPL8 | 6132 | ENSG00000161016 | ribosomal protein L8 | 2 | MNC, Degree | L8 | 8q24.3 |
| 10 | ATP5D | 513 | ENSG00000099624 | ATP synthase F1 subunit delta | 2 | MNC | ATP5F1D, MC5DN5 | 19p13.3 |
| 11 | RPS16 | 6217 | ENSG00000105193 | ribosomal protein S16 | 2 | Degree | S16 | 19q13.2 |

Three identified genes in this study (**Table 2**), including NDUFS3, MT-ND4, and MT-ND6, encode subunits of complex I (CxI). MT-ND6 encodes NADH dehydrogenase 6, part of CxI in the mitochondria. CxI is involved in the first step in the electron transport process, in which the electrons are transferred from NADH to ubiquinone. Electrons are subsequently thrown from ubiquinone through various other enzyme complexes leading to the preparation of energy for ATP generation (26). Ishikawa *et al*. showed that the MT-ND6 mutation resulted in suppression of CxI activity and induction of ROS production, leading to the stimulation of metastasis in breast and lung cancer cells (27). There is no report on the role of MT-ND6 in GC. MT-ND4 encodes NADH dehydrogenase 4, a part of CxI, one of the enzyme complexes involved in OXPHOS. In the whole mitochondrial genome sequencing study in GC, it has been reported that tumor has remarkably more variants in the MT-ND4 region (24). NDUFS3 encodes one of the iron-sulfur protein components of mitochondrial NADH: ubiquinone oxidoreductase (CxI). Recent studies in GC showed the prognostic capabilities of Nicotinamide N-methyltransferase (NNMT). They found that NNMT expression was substantially connected to clinical pathologic stage, tumor size, and lymph node status in GC. Silencing the NNMT gene resulted in the inhibition of proliferation, invasion, and migration of the GC cells. These results represent the potential of NNMT as a beneficial prognostic marker of GC. It has been observed in neuroblastoma that the presence of NNMT could meaningfully reduce the death of SH-SY5Y cells, and the effects of NNMT were correlated with the elevated intracellular ATP content, ATP/ADP ratio, and CxI activity, as well as a decrease in the degradation of the NDUFS3 subunit of CxI (28).

Among the identified genes in this study (**Table 2**), two genes, including COX7A2 and COX6C, encode cytochrome c oxidase (Cytc) subunits. COX7A2 encodes subunit 7A2 of the cytochrome c oxidase, which is the terminal module of the respiratory chain in the mitochondrial and catalyzes the transfer of electrons from the reduced cytochrome c to oxygen. Data extracted from MALDI-MSI experiments have represented that COX7A2 expression is correlated with the survival curve in GC (29). *Et al*. showed that COX7A2 increased expression is a characteristic feature of the intestinal metaplasia in the esophagus (30). COX6C encodes subunit 6c of the COX complex, which catalyzes the final step of the electron transfer chain. Recently, many investigations have reported the unusual level of COX6C in different cancerous and non-cancerous disease conditions such as diabetes, uterine leiomyoma, prostate cancer, melanoma tissues, breast cancer, and follicular thyroid cancer. It has been reported that the overexpression of NDUFA4 leads to substantial upregulation of the COX6C, which subsequently promotes GC cell proliferation and reduces apoptosis in these cells (31).

Most of this study’s identified genes were involved in mitochondrial functions. Recently, many cancer investigations have focused on the mitochondria for further elucidation of the molecular mechanisms of carcinogenesis. Mitochondria have complicated and diverse functions, including cell signaling, modulation of the biosynthetic metabolism, cell death, and their role in energy production. Different studies provided evidence about the potential correlation between the protein alterations and the number of mitochondria, as well as abnormal components and function of the mitochondria with several human cancer progression and prognosis patients, signifying the possible functionality of mitochondria in new anti-tumor therapeutic strategies by targeting the mitochondrial proteins or metabolism (29). However, the characteristics of respiratory and metabolic changes in GC have not still been fully elucidated. The identified genes in this study might be promising diagnostic and prognostic biomarkers for GC. Nonetheless, these results were obtained by bioinformatics analysis and require further clinical validation.

Finally, two ribosomal proteins, RPL8 and RPS16 (**Table 2**), were also identified in the present study. RPL8 encodes ribosomal L8 protein, which is one of the components of the 60S subunit. Based on a bioinformatics analysis-based study, among the genes highly expressed in one of the clusters, few were associated with ribosomal protein-encoding genes such as RPL8 (32). RPS16 encodes ribosomal protein S16. The diseases associated with RPS16 include Diamond-Blackfan Anemia and Descending Colon Cancer. It has been reported that defection in UQCRQ, as a subunit of complex III, can cause mitochondrial dysfunction, which is correlated with the pathogenesis of ulcerative colitis (33).

### Gene Ontology and pathway enrichment analysis of genes

GO analysis is a well-known method for gene and gene products and identification of typical biological aspects of high-throughput genome or transcriptome data, including molecular function (MF), cellular components (CC), and biological process (BP) (34, 35).

The results of GO analysis and pathway enrichment are illustrated in Error! Reference source not found. to **Figure *6***. The predominant (≥50% observed genes) GO terms found for BP are significantly enriched in cellular process, metabolic process, cellular metabolic process, organic substance metabolic process, primary metabolic process, nitrogen compound metabolic process, transport, organonitrogen compound metabolic process, cellular biosynthetic process, organic substance biosynthetic process, cellular nitrogen compound metabolic process, cellular component organization or biogenesis, organonitrogen compound biosynthetic process, cellular nitrogen compound biosynthetic process, and cellular component organization (Error! Reference source not found.). Our results showed that the metabolic processes are the most affected in GC. Recent investigations have focused on the elucidation of the correlation between metabolic reprogramming and the pathogenesis of cancer. Specifically, regulation of the metabolism and cancer investigation is more brought into intense attention with the development of metabolomics. Metabolomics offers comprehensive knowledge of metabolic profiles of certain cancers. It may provide an excellent tool to find biomarkers for prognosis, diagnosis, metastatic surveillance, and therapeutic sensitivity estimate. Various metabolic changes have been identified in GC, including glucose metabolism, amino acid metabolism, lipid metabolism, and nucleotide metabolism. In addition to the mentioned metabolic alterations in GC, changed levels of other metabolites, including creatinine and inositol, have also been reported in GC (36).

**Figure 1.**
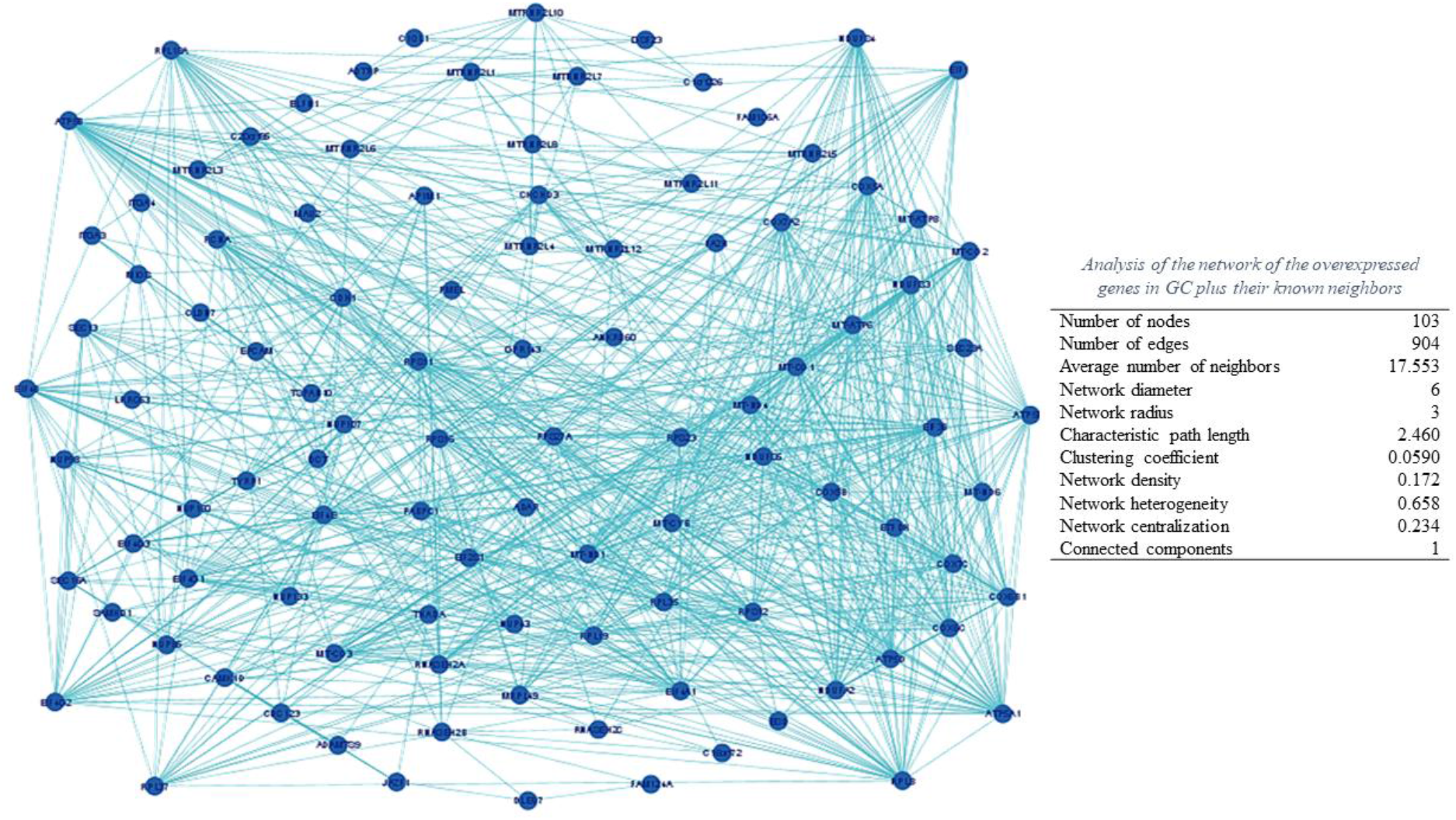
Network of the overexpressed genes in GC plus their known neighbors based on data provided in our previous studies using Cytoscape software.

**Figure 2.**
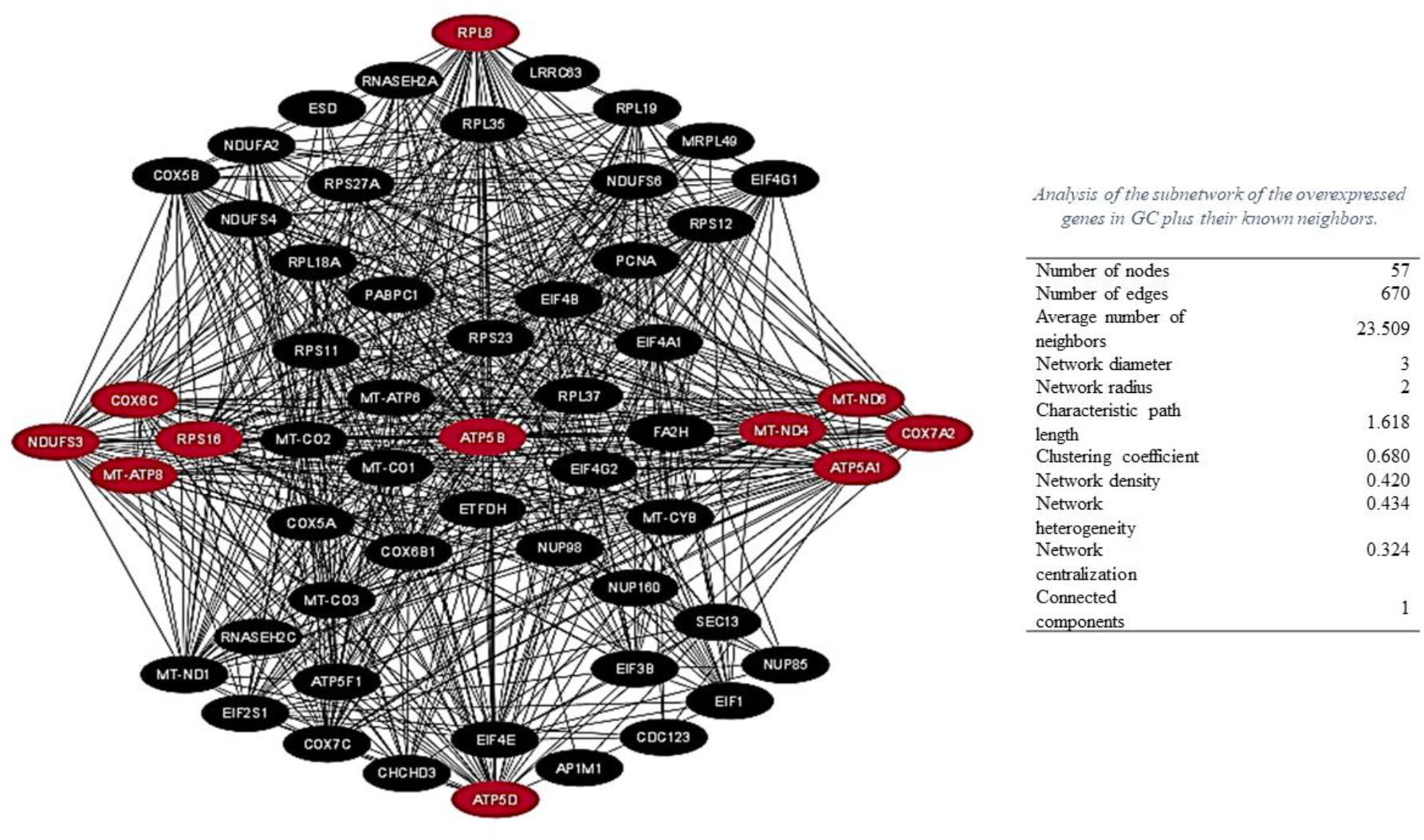
Subnetwork of the overexpressed genes in GC plus their known neighbors based on data provided in our previous studies using the CytoHubba App.

**Figure 3.**
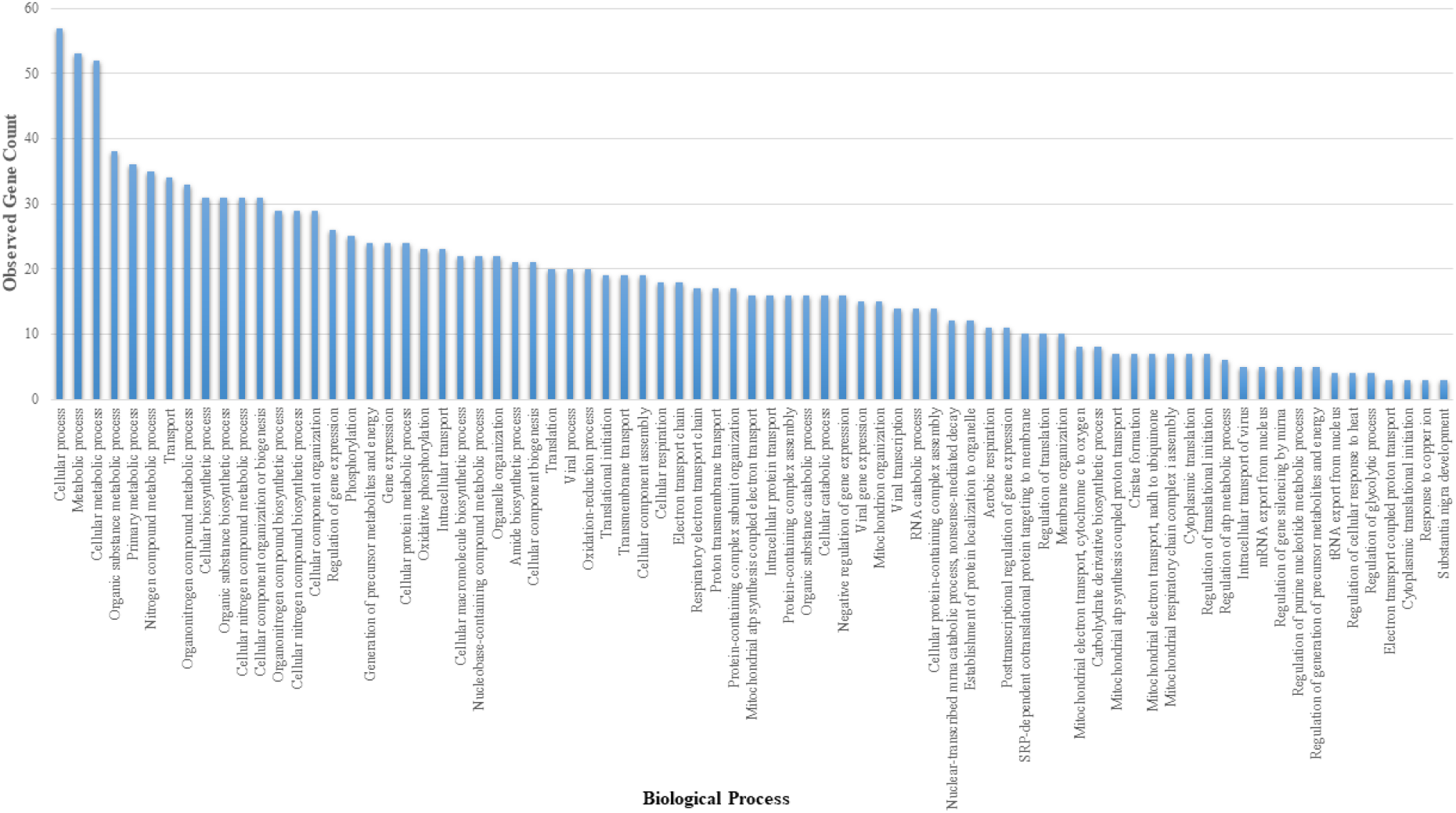
Gene Ontology enrichment analysis (Biological Process) of the determined hub genes in GC based on our previous studies (11-13) data using STRING ver.10 (http://string-db.org).

**Figure 4.**
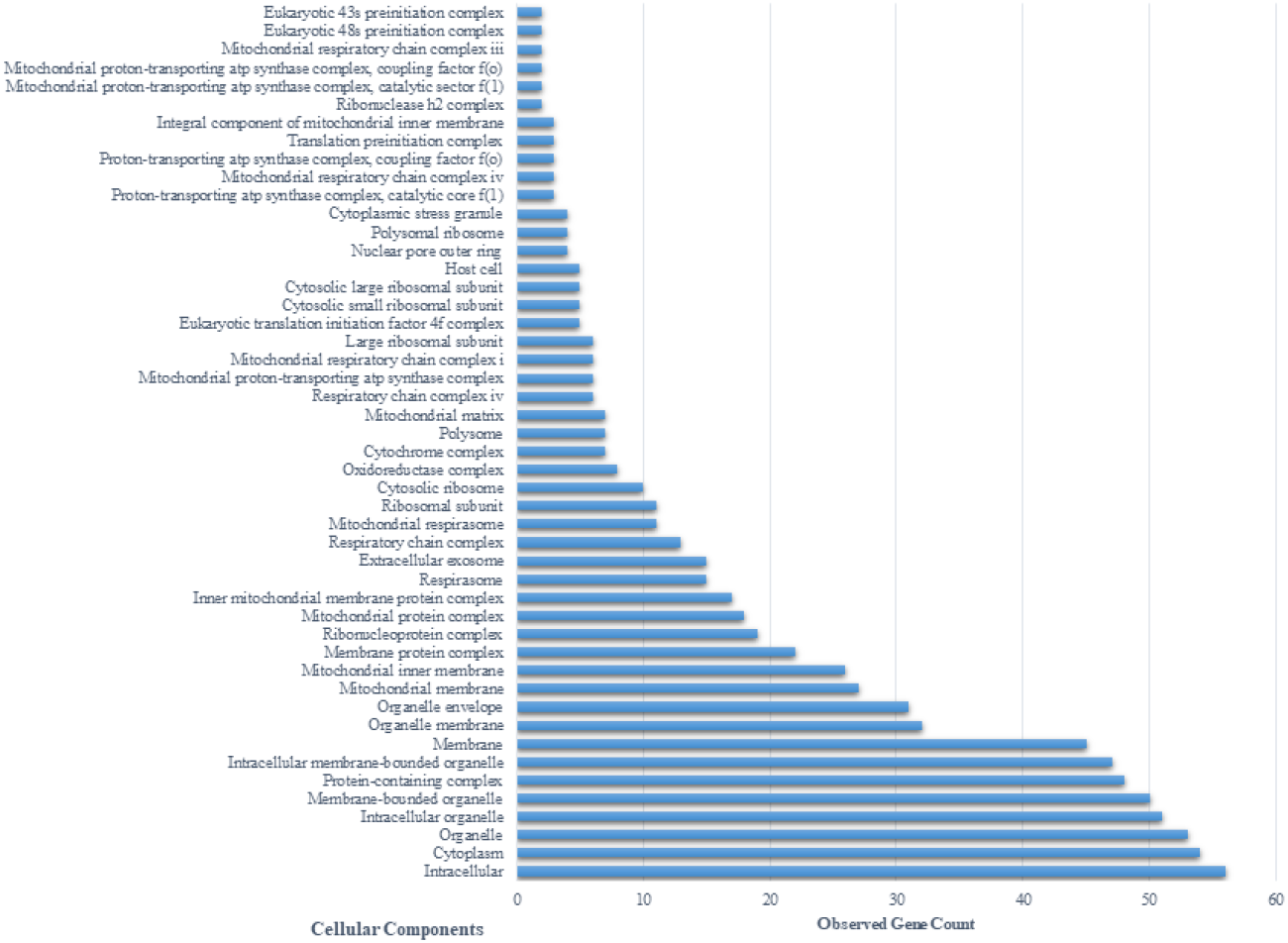
Gene Ontology enrichment analysis (Cellular Component) of the determined hub genes in GC based on our previous studies (11-13) data using STRING ver.10 (http://string-db.org).

**Figure 5.**
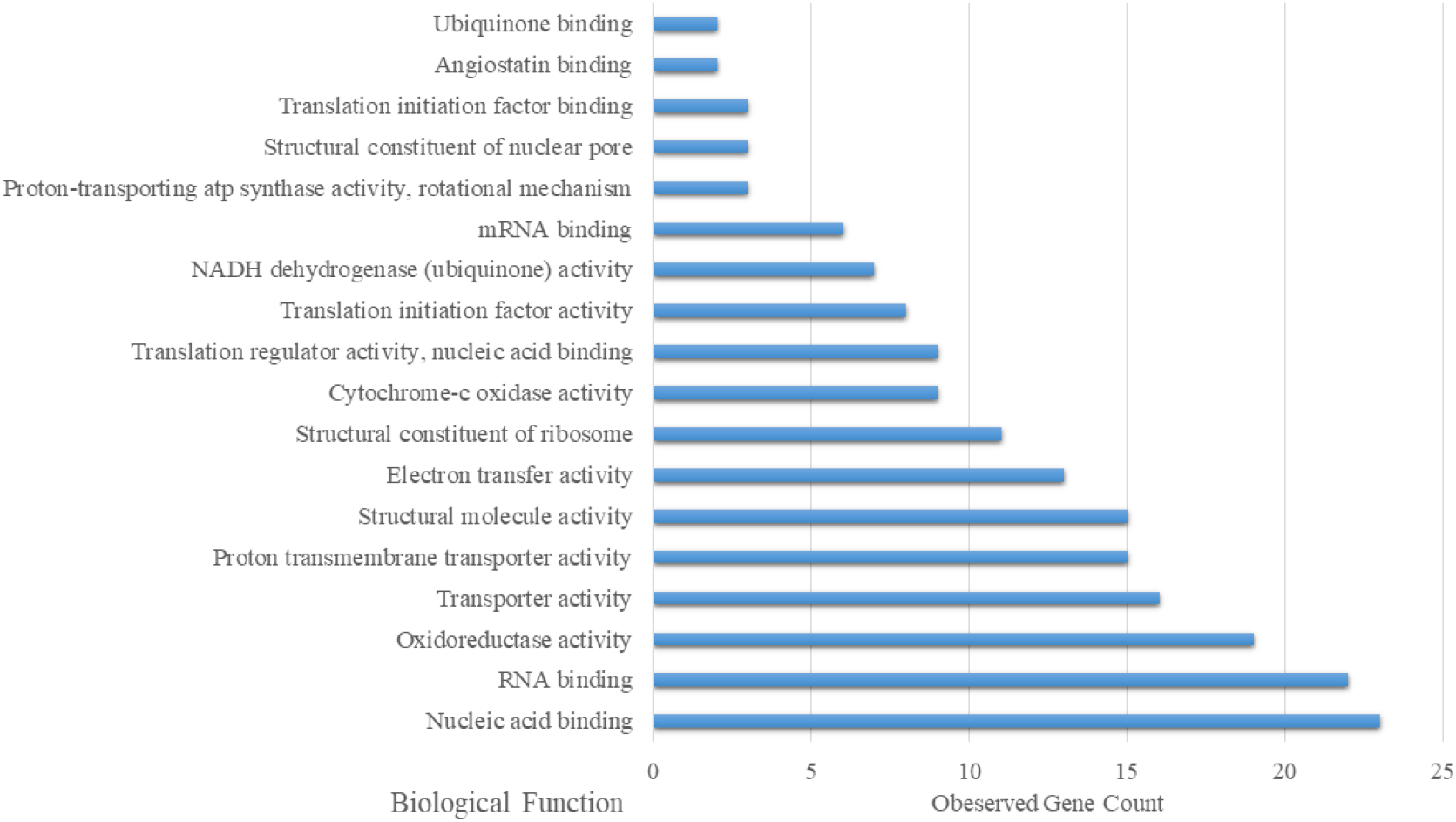
Gene Ontology enrichment analysis (Biological Function) of the determined hub genes in GC based on our previous studies data (11-13) using STRING ver.10 (http://string-db.org).

**Figure 6.**
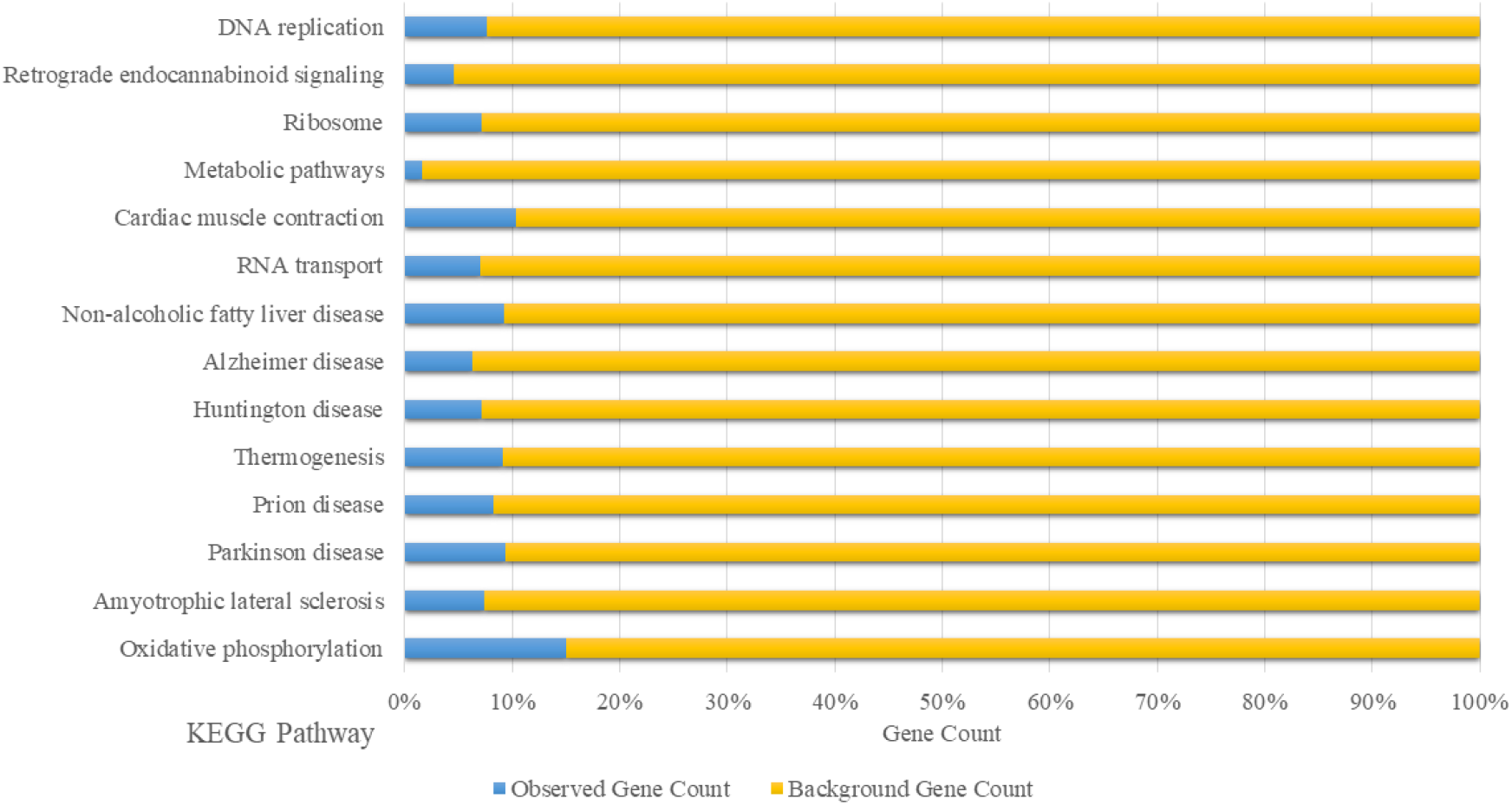
Kyoto Encyclopedia of Genes and Genomes (KEGG) pathways analysis on the hub genes in GC based on our previous studies data (11-13) using STRING ver.10 (http://string-db.org).

The predominant (≥50% observed genes) GO terms found for CC are intracellular, cytoplasm, organelle, intracellular organelle, membrane-bounded organelle, protein-containing complex, intracellular membrane-bounded organelle, membrane, organelle membrane, and organelle envelope (Error! Reference source not found.). The predominant (≥20% observed genes) GO terms found for BF are nucleic acid binding, RNA binding, oxidoreductase activity, transporter activity, proton transmembrane transporter activity, structural molecule activity, and electron transfer activity (Error! Reference source not found.). Our results showed the potential significance of RNAs in GC pathogenesis, confirming recent studies focused on elucidating the role of RNAs in the pathogenesis of this cancer. Using an inclusive assessment of the results from other studies, it has been demonstrated that circular RNAs (circRNAs) regulate the cellular biological behaviors in GC, including epithelial-mesenchymal transition (EMT), proliferation, migration, and invasion. In addition, circRNAs are correlated to the GC characteristics, including tumor stage, differentiation of the tumor, and metastasis. Therefore, circRNAs might be suitable to be used as prognostic or diagnostic biomarkers in GC. In addition, the circRNAs involved in GC pathogenesis can be used as targets in GC treatment (37).

In addition, the GO results in terms of BF showed the significance of oxidoreductase activity, electron transfer activity, and proton transmembrane transporter activity which may all contribute to the mitochondrial function of ATP synthase that has been discussed in previous sections. On the other hand, oxidoreductase activity has been investigated in various cancers, including the breast, liver, gastrointestinal tract, and kidney. It has been shown that Xanthine oxidoreductase (XOR) expression is negatively correlated with poor prognosis in these malignancies (38).

KEGG (http://www.genome.jp/) is a database for allocating certain pathways to groups of DEGs, therefore linking omics data with higher-order functional data (39). Based on KEGG pathway enrichment analysis (**Figure 6**), the prominent (≥40% observed genes) pathways are amyotrophic lateral sclerosis, Parkinson’s disease, Prion disease, Alzheimer’s disease, Metabolic pathways, Oxidative phosphorylation, Thermogenesis, and Huntington’s disease. Most of these pathways are related to neurodegenerative diseases and based on the previous studies (40), SIRT3 is one of the common genes among these pathways. SIRT3 is a member of the sirtuin protein family known as class III histone deacetylases. Recent investigations have focused on SIRT3 due to its function in stress resistance, aging, neurodegenerative disease, and cancer. SIRT3 manages energy requests in situations, including mitochondrial metabolism. The elimination of reactive oxygen species and prevention of cancer cells development or apoptosis are the abilities of SIRT3, which highlights its critical role in cancer and various diseases, including Alzheimer’s disease, amyotrophic lateral sclerosis, Parkinson’s disease, and Huntington’s disease (40).

### Cluster analysis of the network

One of the most important strategies for identifying the functional modules and predicting network biomarkers and complexes of proteins is cluster analysis of the biological networks. In addition, when the results of clustering are visualized, the biological network structure can be displayed. CytoCluster, which is used in this study, includes six clustering algorithms. The selection of the clustering algorithm depends on the user requirements (16). The IPCA algorithm used in this study is density-based, identifying dense subgraphs in protein interaction networks. IPCA determines the weight of an edge by calculating the common neighbors of its connected nodes and calculates each node’s weight through a sum up of the weights of its incident edges. A node needs to have a higher weight to be regarded as a seed. At first, a seed is considered a cluster. IPCA then extends a cluster by adding vertices recursively from its neighbor regarding the nodes’ priority. Adding a node to a cluster depends on two parameters: 1) the probability of its interaction and 2) the shortest path between it and the nodes in the cluster (16). Eleven clusters were obtained in the cluster analysis of the subnetwork. Clusters with ranks 1 to 4 were selected to be discussed in this paper. As shown in **Table 3**, the common pathways among all four cluster ranks include oxidative phosphorylation, Parkinson’s disease, thermogenesis, prion disease, Huntington’s disease, Amyotrophic lateral sclerosis, Alzheimer’s disease, non-alcoholic fatty liver disease, metabolic pathways, cardiac muscle contraction, and retrograde endocannabinoid signaling. However, the Ribosome pathway is not shown only in cluster rank 4, and the DNA replication pathway is only shown in cluster rank 3. These results confirmed the common pathways involved in both neurodegenerative diseases and GC.

**Table 3.**
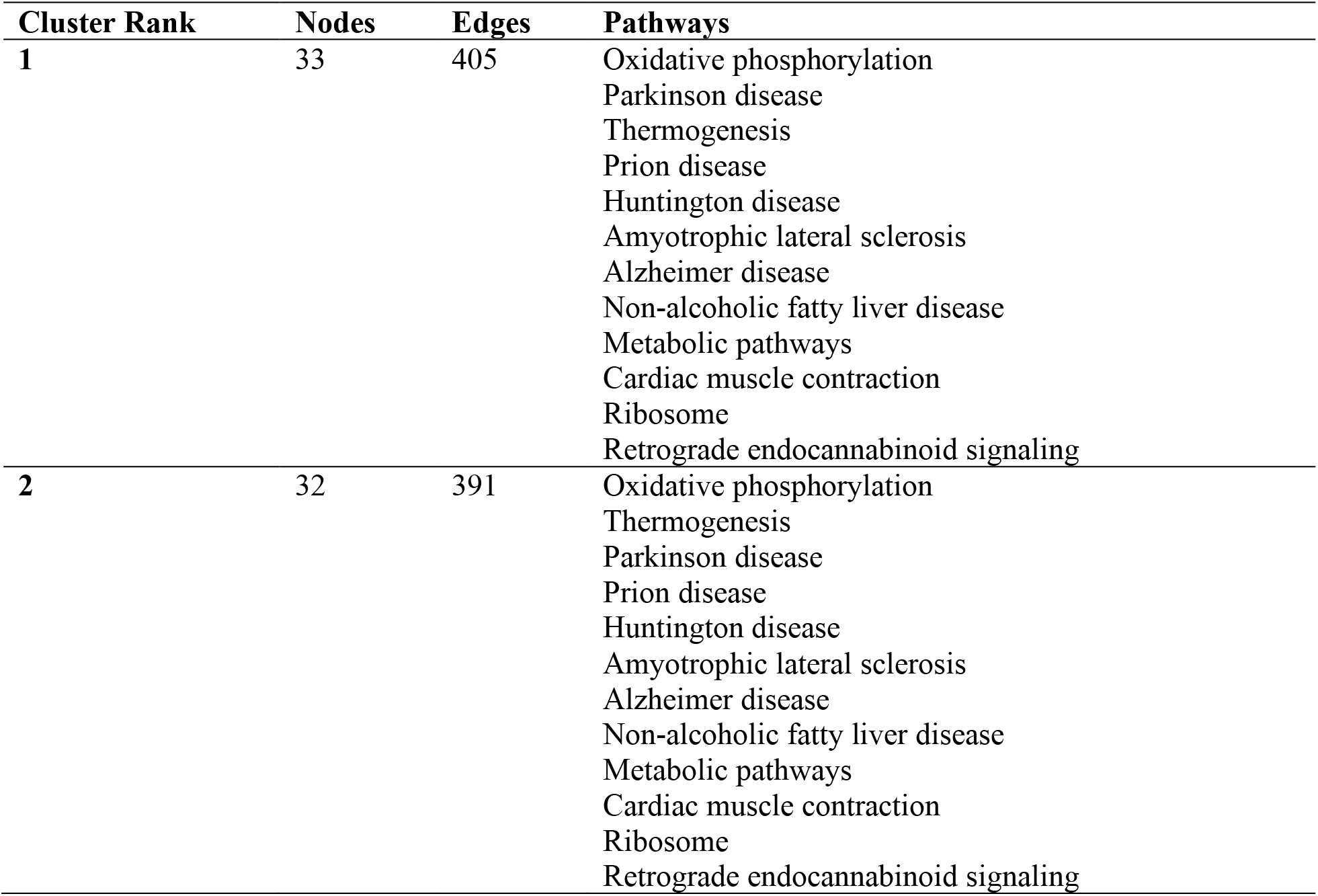
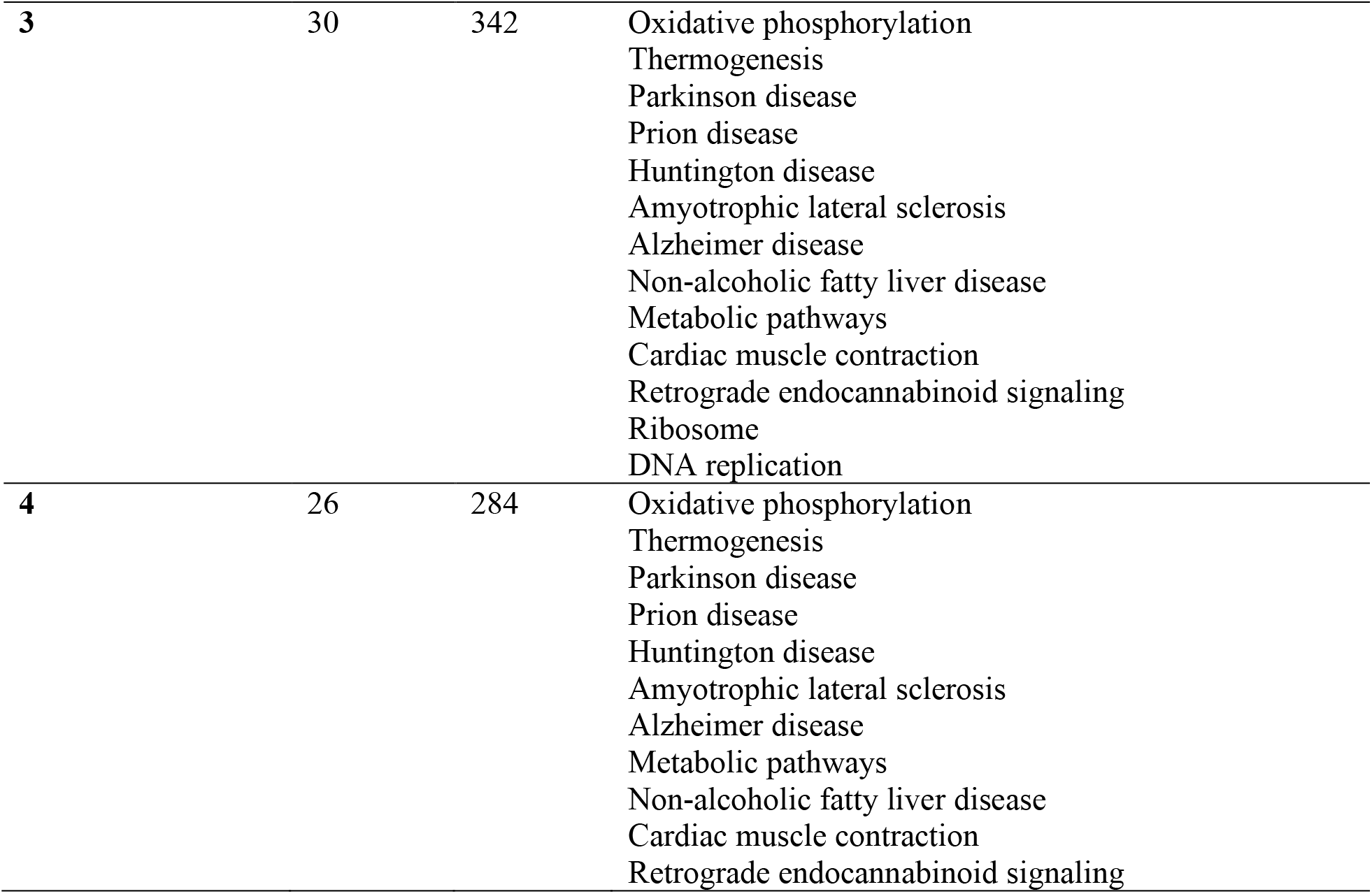
Summary of the clusters (rank 1 to 4) resulted from the cluster analysis of the subnetwork of the overexpressed genes in GC plus their known neighbors based on data provided in our previous studies using the CytoCluster App.

| Cluster Rank | Nodes | Edges | Pathways |
| --- | --- | --- | --- |
| 1 | 33 | 405 | Oxidative phosphorylation<br>Parkinson disease<br>Thermogenesis<br>Prion disease<br>Huntington disease<br>Amyotrophic lateral sclerosis<br>Alzheimer disease<br>Non-alcoholic fatty liver disease<br>Metabolic pathways<br>Cardiac muscle contraction<br>Ribosome<br>Retrograde endocannabinoid signaling |
| 2 | 32 | 391 | Oxidative phosphorylation<br>Thermogenesis<br>Parkinson disease<br>Prion disease<br>Huntington disease<br>Amyotrophic lateral sclerosis<br>Alzheimer disease<br>Non-alcoholic fatty liver disease<br>Metabolic pathways<br>Cardiac muscle contraction<br>Ribosome<br>Retrograde endocannabinoid signaling |
| 3 | 30 | 342 | Oxidative phosphorylation<br>Thermogenesis<br>Parkinson disease<br>Prion disease<br>Huntington disease<br>Amyotrophic lateral sclerosis<br>Alzheimer disease<br>Non-alcoholic fatty liver disease<br>Metabolic pathways<br>Cardiac muscle contraction<br>Retrograde endocannabinoid signaling<br>Ribosome<br>DNA replication |
| 4 | 26 | 284 | Oxidative phosphorylation<br>Thermogenesis<br>Parkinson disease<br>Prion disease<br>Huntington disease<br>Amyotrophic lateral sclerosis<br>Alzheimer disease<br>Metabolic pathways<br>Non-alcoholic fatty liver disease<br>Cardiac muscle contraction<br>Retrograde endocannabinoid signaling |

KEGG pathway analyses showed that the OXPHOS pathway was considerably enriched, and the cluster analysis confirmed it. Recent studies presented that expression levels of several genes, such as NDUFB7, UQCRC2, and UQCRQ in this pathway, were meaningly down-regulated in GC. In addition, studies showed that the enhanced expression of genes in this pathway, such as UQCRQ, NDUFB7, and UQCRC2, positively correlated with a better prognosis (41).

OXPHOS produces energy in most cells which needs several respiratory enzyme complexes and the mitochondrial respiratory chain. The mitochondrial respiratory chain, located in the mitochondria membrane, is composed of four complexes in most organisms. Electron transport is coupled to proton translocation from the matrix of the mitochondria to the intermembrane part, and the enzyme ATP synthase exploits the produced gradient of the proton to generate ATP after phosphorylation of ADP. The switching of cellular energy production from OXPHOS by mitochondria to aerobic glycolysis, named the Warburg effect, happens in several cancer types. One of the major characteristics of fast-growing cancer cells is to maintain an increased glycolysis level providing enough ATP irrespective of the existence of oxygen in the environment (41, 42). However, the importance of this switching for GC development is poorly understood. Recently, *Feichtinger et al*. investigated the expression of OXPHOS complexes in GC using immunohistochemistry. They found that CxI expression was significantly reduced in intestinal-type GC (but not diffuse) GC. Higher complex I and II were seen in larger tumors, and higher expression of complex II and III was observed in higher grades (43).

On the other hand, Su *et al*. found that all of the identified DEPs involved in oxidative phosphorylation were down-regulated. They also reported that GC has substantially changed metabolic processes, including NADH dehydrogenase complex assembly and tricarboxylic acid cycle, consistent with the KEGG analysis results (41). Therefore, the expression status of the OXPHOS complexes in GC needs to be further studied to elucidate the exact role of these complexes in the pathogenesis of GC.

The identified genes in our study (**Table 2**) are mostly involved in mitochondrial functions. Dysfunction of the mitochondrial has been widely described in various cancers. However, the concept of the involvement of both mitochondrial glycolysis and metabolism in cancer cells is controversial. On the other hand, the critical role of the mitochondria in the maintenance of cancer is inevitable. Consequently, there have been several discussions and investigations about the alterations in the function of the mitochondria and their protein expression in various human cancers (33). It has been recently reported that a mutation in the UQCRC2 gene might lead to a deficiency of the mitochondrial complex III, a relatively rare disease. Also, a substantially lower content of UQCRC2 has been reported in breast cancer cells compared to normal cells. Downregulation of this gene has also been reported in glioma and GC. In recent years, many investigations have shown the role of mitochondria in cancer, representing the malfunctioned bioenergetic mitochondria as a symbol of tumorigenesis.

Furthermore, mitochondrial changes have been reported to be correlated with tumor cell migration, invasion, and resistance to chemotherapy (33). These findings suggest that our identified genes might be involved in GC pathogenesis through various pathways. However, the correlation between these genes in regulating mitochondrial function and cancer pathogenesis continues to be unclear and requires to be validated by more research.

### Promoter motif analysis of DEGs

The UFRs (1000 bp) of DEGs were analyzed to identify the conserved motifs and consensus cis-regulatory elements (CREs). The UFRs were extracted using Ensembl BioMarts, a hub for data retrieval across taxonomic space (https://asia.ensembl.org/info/data/biomart/index.html) (44). The MEME Suite web server offers an integrated portal for online finding and assessing motifs indicating features such as protein interaction domains and DNA binding sites. Motifs of transcription factors (TFs) can be compared with motifs in several common motif databases using TOMTOM, an algorithm for scanning the motif databases. Further analysis of TF motifs can be done for putative activities by association with GO terms, including BP, MF, and CC, using GOMO, which is the motif-GO term association tool (17). Accordingly, the extracted UFR sequences were analyzed using TOMTOM (18) to find significant motifs. Subsequently, selected motifs were analyzed using GOMO (19). Ten significant motifs were detected with lengths ranging from 20 to 24nt in length. The results of promoter analysis revealed that KLF16, MAZ, PATZ1, ZNF467, WT1, VEZF1, TBX15, SP1, SP2, and SP3 are among the most common transcription factor families having a binding site in promoters of our hub genes. The GOMO analysis for the motifs discovered by MEME identified several interesting biological functions (**Table 4)**. Gene Ontology indicated that these motifs participated in potassium ion transport, negative regulation of neuron apoptosis, neuron fate commitment, and inner ear morphogenesis. However, involvement in the negative regulation of signal transduction was the most common biological process among all motifs. Therefore, this biological process may play an important role in GC pathogenesis. In addition, the predominant GO terms found for CC were significantly enriched in transcription factor complex and dendrite. Moreover, these motifs involved molecular functions, including chromatin binding, protein homodimerization activity, protein heterodimerization activity, potassium ion binding, and transcription activator activity. Therefore, these biological processes play an important role in the pathogenesis of GC.

**Table 4.**
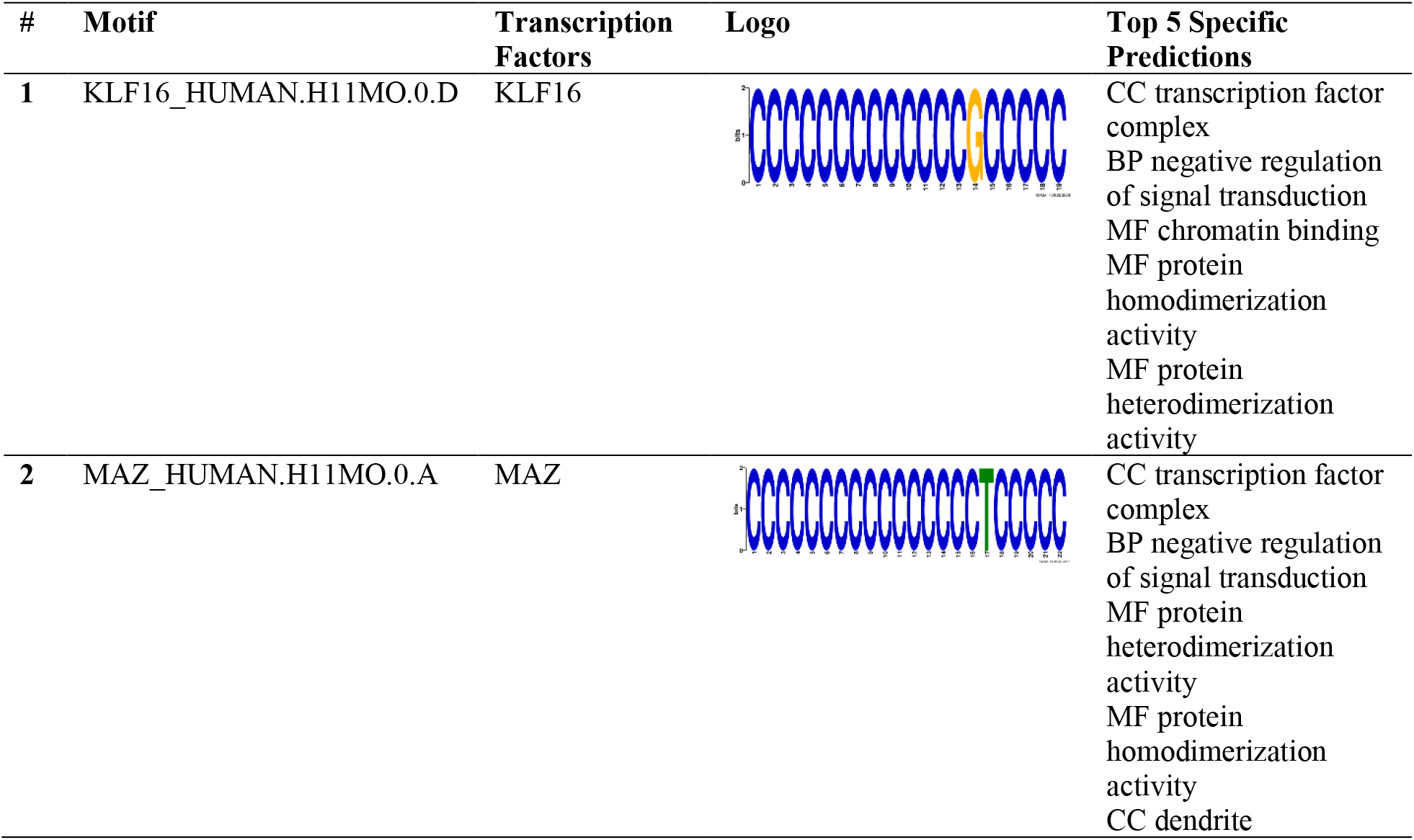
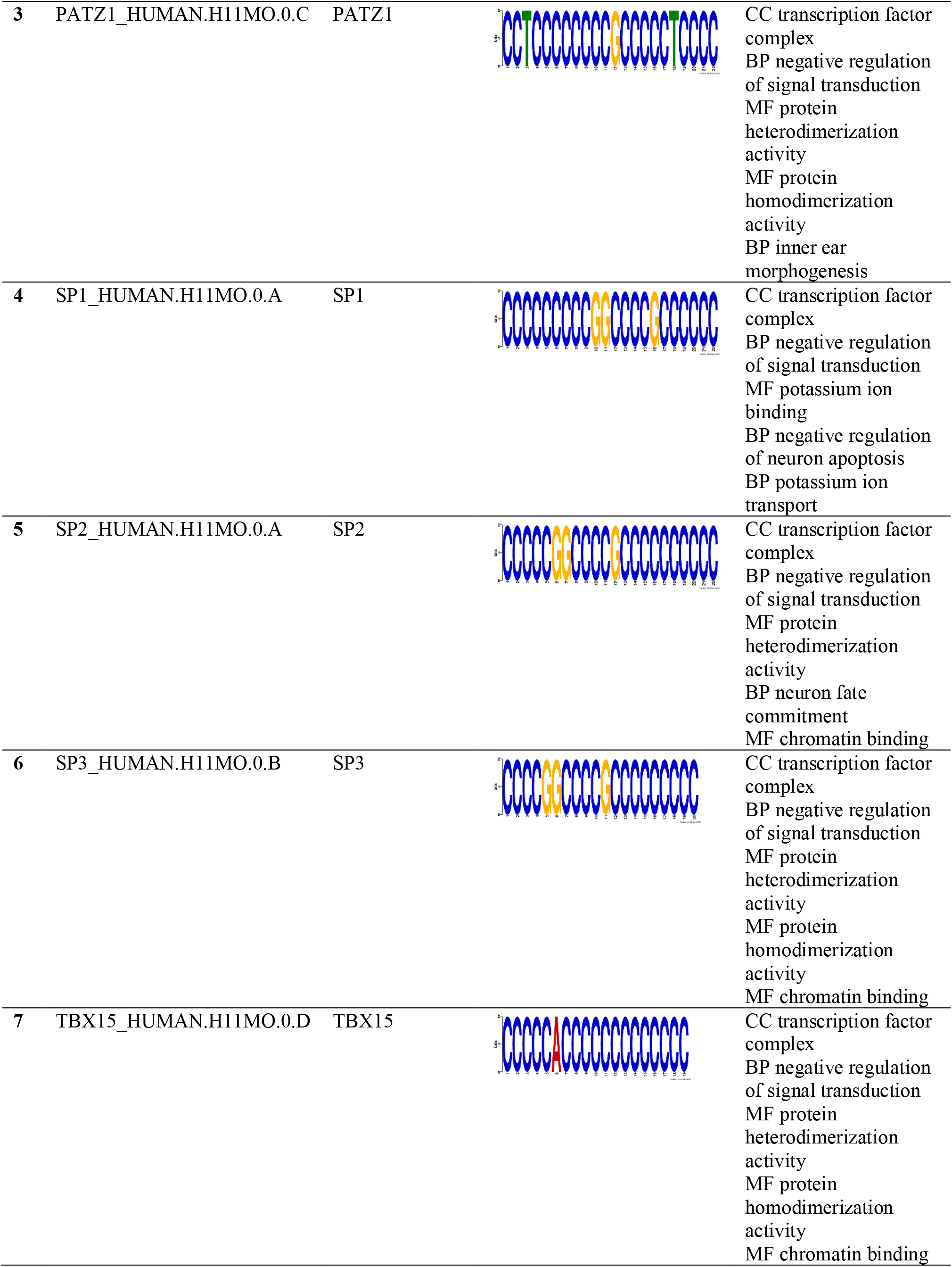
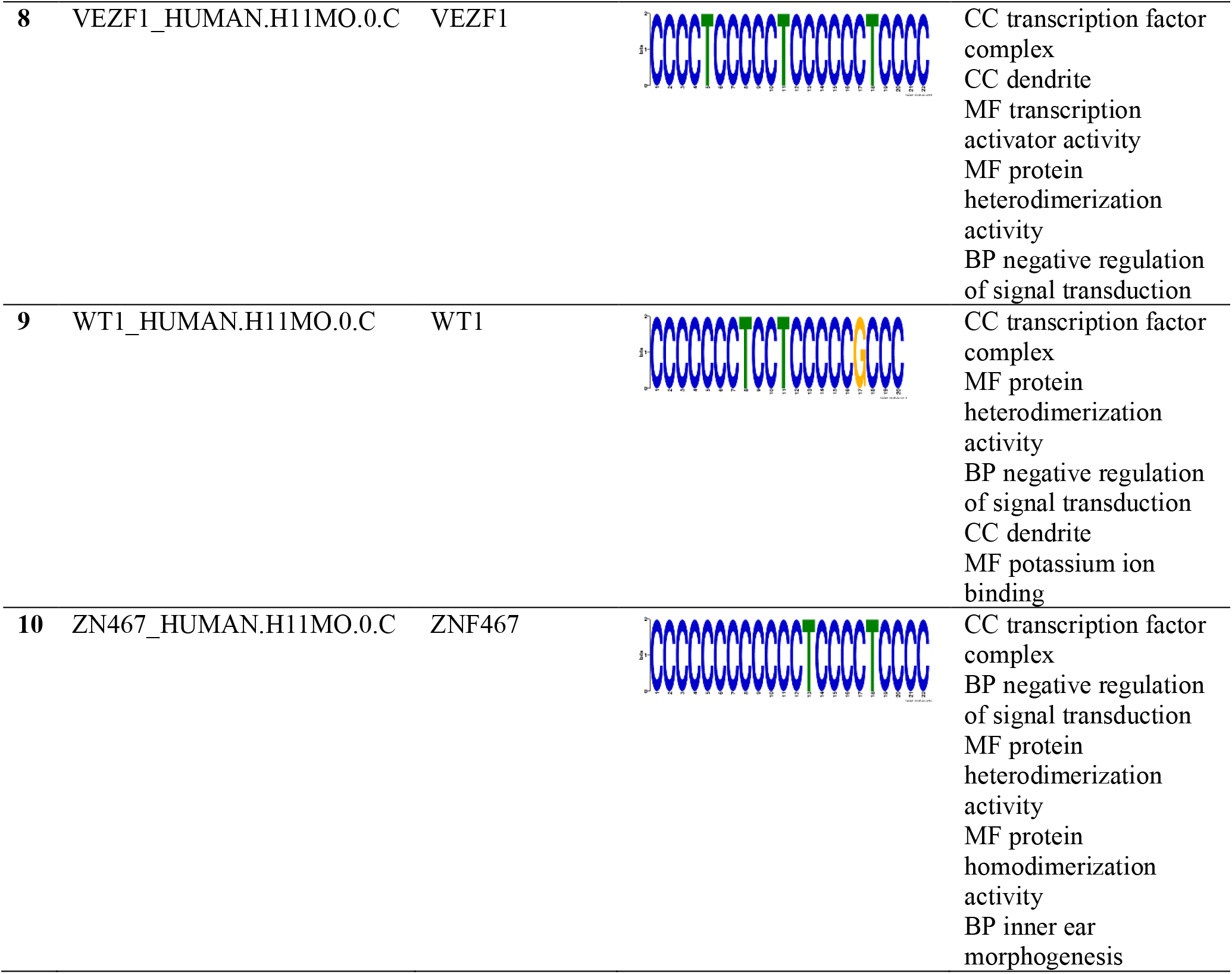
the conserved motifs found in promoters of DEGs by MEME analysis.

| # | Motif | Transcription Factors | Logo | Top 5 Specific Predictions |
| --- | --- | --- | --- | --- |
| 1 | KLF16_HUMAN.H11MO.0.D | KLF16 |  | CC transcription factor complex<br>BP negative regulation of signal transduction<br>MF chromatin binding<br>MF protein homodimerization activity<br>MF protein heterodimerization activity |
| 2 | MAZ_HUMAN.H11MO.0.A | MAZ |  | CC transcription factor complex<br>BP negative regulation of signal transduction<br>MF protein heterodimerization activity<br>MF protein homodimerization activity<br>CC dendrite |
| 3 | PATZ1_HUMAN.H11MO.0.C | PATZ1 |  | CC transcription factor complex<br>BP negative regulation of signal transduction<br>MF protein heterodimerization activity<br>MF protein homodimerization activity<br>BP inner ear morphogenesis |
| 4 | SP1_HUMAN.H11MO.0.A | SP1 |  | CC transcription factor complex<br>BP negative regulation of signal transduction<br>MF potassium ion binding<br>BP negative regulation of neuron apoptosis<br>BP potassium ion transport |
| 5 | SP2_HUMAN.H11MO.0.A | SP2 |  | CC transcription factor complex<br>BP negative regulation of signal transduction<br>MF protein heterodimerization activity<br>BP neuron fate commitment<br>MF chromatin binding |
| 6 | SP3_HUMAN.H11MO.0.B | SP3 |  | CC transcription factor complex<br>BP negative regulation of signal transduction<br>MF protein heterodimerization activity<br>MF protein homodimerization activity<br>MF chromatin binding |
| 7 | TBX15_HUMAN.H11MO.0.D | TBX15 |  | CC transcription factor complex<br>BP negative regulation of signal transduction<br>MF protein heterodimerization activity<br>MF protein homodimerization activity<br>MF chromatin binding |
| 8 | VEZF1_HUMAN.H11MO.0.C | VEZF1 |  | CC transcription factor complex<br>CC dendrite<br>MF transcription activator activity<br>MF protein heterodimerization activity<br>BP negative regulation of signal transduction |
| 9 | WT1_HUMAN.H11MO.0.C | WT1 |  | CC transcription factor complex<br>MF protein heterodimerization activity<br>BP negative regulation of signal transduction<br>CC dendrite<br>MF potassium ion binding |
| 10 | ZN467_HUMAN.H11MO.0.C | ZNF467 |  | CC transcription factor complex<br>BP negative regulation of signal transduction<br>MF protein heterodimerization activity<br>MF protein homodimerization activity<br>BP inner ear morphogenesis |

Although our study suggested the probable relevance between these identified genes/CREs and GC pathogenesis, little is known about the relevance of these genes/CREs to GC. Moreover, clinical validation and long-term follow-up data for GC patients with a large sample size are required for investigating these genes as prognostic or diagnostic biomarkers. If these genes’ diagnostic/prognostic importance is validated, they could be introduced as factors to define the treatment modality for GC. Research on targeted changes in the expression of these genes can offer novel therapeutic modalities.

## Conclusion

Eleven hub genes in GC were identified in the current study, mostly involved in mitochondrial functions. The GO and pathway analysis showed that the metabolic processes are most affected in GC. These analyses also confirmed the potential significance of mitochondrial role in GC pathogenesis. Although many investigations have been focused on the mitochondria further to elucidate the potential contribution of it to carcinogenesis, however, the characteristics of respiratory and metabolic changes in GC have not been fully elucidated. In addition, KEGG analysis and cluster analysis results showed that most of the affected pathways in GC were the pathways also involved in neurodegenerative diseases. KEGG pathway analyses also showed that the OXPHOS pathway was considerably enriched, and the cluster analysis confirmed it. Also, the promoter analysis results showed that negative signal transduction regulation might play an important role in GC pathogenesis. The results of this study might open new insights into GC pathogenesis. In addition, the identified genes in this study might be promising diagnostic and prognostic biomarkers or therapeutic targets for GC. Nonetheless, these results were obtained by bioinformatics analysis and require further clinical validation.

## Conflict of interest

The authors declare no conflict of interest.

## Acknowledgment

We would like to acknowledge the Iran University of Medical Sciences and Nuclear Science and Technology Research Institute for their support. We also thank Iman Tavassoly (MD, Ph.D.) for editing the paper.

## Funding/support

This study received no specific grant from any funding agency in the public, commercial, or not-for-profit sectors.

## Authors’ Contribution

Study concept and design: NMD. and AG ; Acquisition of data: NMD and HM; Drafting of the manuscript: NMD and AG; Critical revision of the manuscript for important intellectual content: NMD, AG and HM; Study supervision: NMD and AG.

## Notes

### Competing Interest Statement

The authors have declared no competing interest.

## References

1. Sung H, Ferlay J, Siegel RL, Laversanne M, Soerjomataram I, Jemal A, et al. Global Cancer Statistics 2020: GLOBOCAN Estimates of Incidence and Mortality Worldwide for 36 Cancers in 185 Countries. CA: A Cancer Journal for Clinicians. 2021;71(3):209–49.

2. Smyth EC, Nilsson M, Grabsch HI, van Grieken NC, Lordick F. Gastric cancer. The Lancet. 2020;396(10251):635–48.

3. Vogelstein B, Kinzler KW. Cancer genes and the pathways they control. Nature medicine. 2004;10(8):789–99.

4. Boussioutas A, Li H, Liu J, Waring P, Lade S, Holloway AJ, et al. Distinctive patterns of gene expression in premalignant gastric mucosa and gastric cancer. Cancer research. 2003;63(10):2569–77.

5. Nagini S. Carcinoma of the stomach: A review of epidemiology, pathogenesis, molecular genetics and chemoprevention. World journal of gastrointestinal oncology. 2012;4(7):156.

6. Kulasingam V, Diamandis EP. Strategies for discovering novel cancer biomarkers through utilization of emerging technologies. Nature clinical practice Oncology. 2008;5(10):588–99.

7. Tavassoly I, Hu Y, Zhao S, Mariottini C, Boran A, Chen Y, et al. Genomic signatures defining responsiveness to allopurinol and combination therapy for lung cancer identified by systems therapeutics analyses. Molecular Oncology. 2019;13(8):1725–43.

8. Mehrpooya A, Saberi-Movahed F, Azizizadeh N, Rezaei-Ravari M, Saberi-Movahed F, Eftekhari M, et al. High dimensionality reduction by matrix factorization for systems pharmacology. Briefings in Bioinformatics. 2021;23(1).

9. Dorvash M, Farahmandnia M, Tavassoly I. A Systems Biology Roadmap to Decode mTOR Control System in Cancer. Interdisciplinary Sciences: Computational Life Sciences. 2020;12(1):1–11.

10. Tavassoly I, Parmar J, Shajahan-Haq A, Clarke R, Baumann W, Tyson J. Dynamic Modeling of the Interaction Between Autophagy and Apoptosis in Mammalian Cells. CPT: Pharmacometrics & Systems Pharmacology. 2015;4(4):263–72.

11. Mottaghi-Dastjerdi N, Soltany-Rezaee-Rad M, Sepehrizadeh Z, Roshandel G, Ebrahimifard F, Setayesh N. Genome expression analysis by suppression subtractive hybridization identified overexpression of Humanin, a target gene in gastric cancer chemoresistance. DARU Journal of Pharmaceutical Sciences. 2014;22(1):1–7.

12. Mottaghi-Dastjerdi N, Soltany-Rezaee-Rad M, Sepehrizadeh Z, Roshandel G, Ebrahimifard F, Setayesh N. Identification of novel genes involved in gastric carcinogenesis by suppression subtractive hybridization. Human & experimental toxicology. 2015;34(1):3–11.

13. Mottaghi-Dastjerdi N, Soltany-Rezaee-Rad M, Sepehrizadeh Z, Roshandel G, Ebrahimifard F, Setayesh N. Gene expression profiling revealed overexpression of vesicle amine transport protein-1 (VAT-1) as a potential oncogene in gastric cancer. 2016.

14. Szklarczyk D, Franceschini A, Wyder S, Forslund K, Heller D, Huerta-Cepas J, et al. STRING v10: protein–protein interaction networks, integrated over the tree of life. Nucleic acids research. 2015;43(D1):D447–D52.

15. Smoot ME, Ono K, Ruscheinski J, Wang P-L, Ideker T. Cytoscape 2.8: new features for data integration and network visualization. Bioinformatics. 2010;27(3):431–2.

16. Li M, Li D, Tang Y, Wu F, Wang J. CytoCluster: a cytoscape plugin for cluster analysis and visualization of biological networks. International journal of molecular sciences. 2017;18(9):1880.

17. Bailey TL, Boden M, Buske FA, Frith M, Grant CE, Clementi L, et al. MEME SUITE: tools for motif discovery and searching. Nucleic Acids Res. 2009;37(Web Server issue):W202–8.

18. Gupta S, Stamatoyannopoulos JA, Bailey TL, Noble WS. Quantifying similarity between motifs. Genome Biology. 2007;8(2):R24.

19. Buske FA, Bodén M, Bauer DC, Bailey TL. Assigning roles to DNA regulatory motifs using comparative genomics. Bioinformatics. 2010;26(7):860–6.

20. Qi B, Han M. Microbial Siderophore Enterobactin Promotes Mitochondrial Iron Uptake and Development of the Host via Interaction with ATP Synthase. Cell. 2018;175(2):571-82.e11.

21. Liu F, Zhang Y, Men T, Jiang X, Yang C, Li H, et al. Quantitative proteomic analysis of gastric cancer tissue reveals novel proteins in platelet-derived growth factor b signaling pathway. Oncotarget. 2017;8(13):22059.

22. Wang X, Chang X, He C, Fan Z, Yu Z, Yu B, et al. ATP5B promotes the metastasis and growth of gastric cancer by activating the FAK/AKT/MMP2 pathway. The FASEB Journal. 2021;35(4):e20649.

23. Dautant A, Meier T, Hahn A, Tribouillard-Tanvier D, Di Rago J-P, Kucharczyk R. ATP synthase diseases of mitochondrial genetic origin. Frontiers in physiology. 2018;9:329.

24. Cavalcante GC, Marinho ANR, Anaissi AK, Vinasco-Sandoval T, Ribeiro-dos-Santos A, Vidal AF, et al. Whole mitochondrial genome sequencing highlights mitochondrial impact in gastric cancer. Scientific Reports. 2019;9(1):15716.

25. Wei J, Xie Q, Liu X, Wan C, Wu W, Fang K, et al. Identification the prognostic value of glutathione peroxidases expression levels in acute myeloid leukemia. Annals of Translational Medicine. 2020;8(11).

26. Yuan Y, Wang W, Li H, Yu Y, Tao J, Huang S, et al. Nonsense and missense mutation of mitochondrial ND6 gene promotes cell migration and invasion in human lung adenocarcinoma. BMC cancer. 2015;15(1):1–10.

27. Lin Y-H, Lim S-N, Chen C-Y, Chi H-C, Yeh C-T, Lin W-R. Functional Role of Mitochondrial DNA in Cancer Progression. International journal of molecular sciences. 2022;23(3):1659.

28. Lu X, Long H. Nicotinamide N-methyltransferase as a potential marker for cancer. Neoplasma. 2018;65(5):656–63.

29. Balluff B, Elsner M, Kowarsch A, Rauser S, Meding S, Schuhmacher C, et al. Classification of HER2/neu Status in Gastric Cancer Using a Breast-Cancer Derived Proteome Classifier. Journal of Proteome Research. 2010;9(12):6317–22.

30. Elsner M, Rauser S, Maier S, Schöne C, Balluff B, Meding S, et al. MALDI imaging mass spectrometry reveals COX7A2, TAGLN2 and S100-A10 as novel prognostic markers in Barrett’s adenocarcinoma. Journal of Proteomics. 2012;75(15):4693–704.

31. Tian B-X, Sun W, Wang S-H, Liu P-J, Wang Y-C. Differential expression and clinical significance of COX6C in human diseases. American Journal of Translational Research. 2021;13(1):1.

32. Kwon CH, Park HJ, Choi YR, Kim A, Kim HW, Choi JH, et al. PSMB8 and PBK as potential gastric cancer subtype-specific biomarkers associated with prognosis. Oncotarget. 2016;7(16):21454.

33. Zong WX, Rabinowitz JD, White E. Mitochondria and Cancer. Mol Cell. 2016;61(5):667–76.

34. Consortium GO. The gene ontology (GO) project in 2006. Nucleic acids research. 2006;34(Suppl_1):D322–D6.

35. Tavassoly I, Goldfarb J, Iyengar R. Systems biology primer: the basic methods and approaches. Essays in Biochemistry. 2018;62(4):487–500.

36. Xiao S, Zhou L. Gastric cancer: metabolic and metabolomics perspectives. International Journal of Oncology. 2017;51(1):5–17.

37. Fang X, Wen J, Sun M, Yuan Y, Xu Q. CircRNAs and its relationship with gastric cancer. Journal of Cancer. 2019;10(24):6105.

38. Battelli MG, Polito L, Bortolotti M, Bolognesi A. Xanthine oxidoreductase in cancer: more than a differentiation marker. Cancer Medicine. 2016;5(3):546–57.

39. Kanehisa M, Goto S. KEGG: kyoto encyclopedia of genes and genomes. Nucleic acids research. 2000;28(1):27–30.

40. Ansari A, Rahman MS, Saha SK, Saikot FK, Deep A, Kim K-H. Function of the SIRT3 mitochondrial deacetylase in cellular physiology, cancer, and neurodegenerative disease. Aging Cell. 2017;16(1):4–16.

41. Su F, Zhou F-f, Zhang T, Wang D-w, Zhao D, Hou X-m, et al. Quantitative proteomics identified 3 oxidative phosphorylation genes with clinical prognostic significance in gastric cancer. Journal of Cellular and Molecular Medicine. 2020;24(18):10842–54.

42. Tavassoly I. Dynamics of Cell Fate Decision Mediated by the Interplay of Autophagy and Apoptosis in Cancer Cells: Mathematical Modeling and Experimental Observations: Springer; 2015.

43. Feichtinger RG, Neureiter D, Skaria T, Wessler S, Cover TL, Mayr JA, et al. Oxidative Phosphorylation System in Gastric Carcinomas and Gastritis. Oxidative Medicine and Cellular Longevity. 2017;2017:1320241.

44. Kinsella RJ, Kähäri A, Haider S, Zamora J, Proctor G, Spudich G, et al. Ensembl BioMarts: a hub for data retrieval across taxonomic space. Database. 2011;2011.

